# Loss of nsp14-exonuclease activity impairs the replication, proofreading, fitness and pathogenesis of SARS-CoV-2

**DOI:** 10.64898/2026.01.12.698941

**Authors:** Jordan Anderson-Daniels, Meghan V. Diefenbacher, Boyd L. Yount, Rita M. Meganck, Longping V. Tse, Kaitlyn N. Burke, Hector A. Miranda, D. Trevor Scobey, Xiaotao Lu, Laura Stevens, Kenneth H. Dinnon, Nathaniel S. Chapman, Camryn Pajon, John M. Powers, Cameron Nguyen, Rachel L Graham, Nicholas S. Heaton, Ralph S. Baric, Mark R. Denison, Timothy P. Sheahan

## Abstract

Coronaviruses (CoVs) replicate their RNA genomes with increased fidelity than other RNA viruses, a mechanism mediated by the proofreading and recombination activities of the exoribonuclease domain of replicase nonstructural protein 14 (nsp14-ExoN). Both murine hepatitis virus (MHV) and SARS-CoV tolerate nsp14-ExoN loss-of-function mutations (ExoN-) (D90A and E92A) but have impaired replication fidelity and pathogenesis yet identical substitutions in MERS-CoV and SARS-CoV-2 have been reported to be lethal. Here, we report a saturation mutagenesis approach facilitating the recovery and analysis of several constellations of SARS-CoV-2 nsp14 ExoN-inactivating, loss-of-function substitutions, including the canonical D90A and E92A. Biochemical assays with purified WT or ExoN- nsp10-14 fusion proteins confirmed that active site substitutions abolished ExoN activity (ExoN-). SARS-CoV-2 ExoN- viruses had impaired replication, RNA synthesis, and recombination, as well as decreased replication fidelity and loss of fitness *in vitro*. ExoN- viruses were significantly attenuated for replication in human primary airway epithelial cells and were attenuated for replication and pathogenesis in WT mice as well as the highly susceptible K18 transgenic mice. In the absence of interferon signaling *in vivo*, SARS-CoV and SARS-CoV-2 ExoN- viral replication could be nearly fully restored. These results demonstrate that SARS-CoV-2 ExoN- are viable but highly impaired for replication, fitness, and fidelity *in vitro* as well as innate immune antagonism and pathogenesis *in vivo*. Collectively, our results solidify the multiple critical roles for nsp14-ExoN across CoV genera and establish new approaches for rescue and analysis of loss-of-function substitutions for studies of CoV replication, pathogenesis, and evolution.

## Introduction

Coronaviruses (CoVs) are etiologic agents of both endemic and pandemic human disease. Three novel CoV have emerged in the past 20 years including severe acute respiratory syndrome CoV (SARS-CoV), Middle East respiratory syndrome CoV (MERS-CoV) and SARS-CoV-2, the causative agent of pandemic COVID-19. All human CoV (HCoV) are thought to have originated from zoonoses (1–5). Given the great diversity of CoVs in animal reservoir species, future spillover events may give rise to novel human diseases. Thus, understanding the mechanisms by which CoVs replicate and evolve is critical for the understanding of emergence potential and the treatment and prevention of current and future emerging CoV infections. Much of the success in the development of COVID-19 antiviral therapies has resulted from the many viral, biochemical, and structural studies of the CoV replication transcription complex (RTC). The CoV RTC is assembled from enzymatic and non-enzymatic co-factor proteins including the non-structural proteins 7-16 (nsp7-16) where the nsp12 RNA-dependent RNA polymerase (nsp12-RdRP) along with cofactors nsp7 and nsp8 drive genome replication, sub-genomic mRNA synthesis, and initial guanylyl-transferase activity to initiate RNA capping (5–7). CoVs and other members of the order *Nidovirales* encode a 3’ to 5’ exoribonuclease (nsp14-ExoN) that regulates replication fidelity through RNA-dependent RNA proofreading functions during RNA synthesis (8–11). RNA proofreading has been proposed as a critical determinant of nidovirus evolution, pathogenesis and the maintenance of their large RNA genomes while using an otherwise low-fidelity RdRp (9, 12–14). ExoN is a shared motif in all nidoviruses with genomes greater than 20kb, and recent transcriptome mining has identified nidoviruses with genomes as large as 64 kb, twice that of SARS-CoV-2 (9, 15–17). Thus, nsp14-ExoN is a central determinant of CoV replication fidelity and evolutionary potential.

The first evidence that nidovirus encoded an ExoN domain was identified through bioinformatic approaches with SARS-CoV nsp14. SARS-CoV nsp14 ExoN activity was subsequently confirmed in biochemical experiments (10, 18). Nsp14-ExoN is a member of the dnaQ-like DE-D-DH family of RNA and DNA exonucleases (19). The CoV ExoN contains three metal coordinating active site motifs (i.e. motif I-Asp-X-Glu (DE), motif II-Glu (E), and motif III-Asp-His (DH)) which give the DE-D-DH family its name although CoVs have a unique substitution in motif II making their active site residue set DE-E-DH (10). ExoN hydrolyses ssRNA and dsRNA in a 3’-5’ direction and excises 3’ single-nucleotide mismatches in template RNAs, activities that require binding of the nonenzymatic nsp10 cofactor (10, 20–22). Alanine substitutions of MHV and SARS-CoV motif I (DE to AA) (nsp14-ExoN-) result in significant defects in RNA replication fidelity and proofreading, thus are “loss of function” (LOF) mutations (12, 13). Importantly, ExoN inactivating mutations attenuate replication and pathogenesis in mice but the underlying mechanisms are not completely understood (23).

While ExoN active site substitutions were tolerated in MHV and SARS-CoV, similar substitutions in ExoN for other CoVs including HCoV-229E, MERS-CoV, and SARS-CoV-2 have not been viable despite these viruses sharing completely conserved active site domain motifs and highly conserved structures with SARS-CoV (10, 24, 25). Here, we report the recovery and analysis of SARS-CoV-2 viruses with nsp14-ExoN LOF substitutions (ExoN-) in the motif-I active site. Using a saturation mutagenesis approach, a highly permissive engineered Vero E6 cell line and low temperature, we recovered a series of viable motif-I substitutions in SARS-CoV-2 ExoN. These experimental conditions also facilitated the recovery of a SARS-CoV-2 with classical motif-1 DE-AA substitutions. Biochemical assays confirmed that mutations found in viable viruses abolished ExoN enzymatic activity. SARS-CoV-2 ExoN- exhibited impaired replication, altered RNA synthesis, loss of competitive fitness, increased sensitivity to nucleoside analogues, decreased fidelity, and altered recombination patterns in the highly permissive and notably interferon (IFN) incompetent Vero cells. SARS-CoV-2 ExoN- viruses were highly attenuated for replication in primary human airway epithelial cells and in WT and highly permissive K18 ACE2 transgenic mice. The attenuation of SARS-CoV-2 ExoN- replication could be partially reversed in mice lacking type I and type III IFN signaling. Altogether, these data indicate that *in vivo* attenuation of ExoN- viruses is not solely driven by general defects in fidelity and replicative capacity but rather highlight the role of an ExoN-mediated innate immune antagonism that is important for *in vivo* replication and pathogenic potential. Thus, our results with SARS-CoV-2 further establish that LOF nsp14-ExoN- mutations result in highly conserved biochemical, virological, and pathogenic defects across divergent CoVs. Finally, these studies demonstrate the importance of nsp14-ExoN as a key regulator replication, pathogenesis and immune evasion making it an ideal target for broadly active therapeutic interventions.

## Results

### SARS-CoV-2 nsp14-ExoN mutants are viable but attenuated for replication in vitro

The SARS-CoV-2 nsp14-ExoN has five active site residues (D90, E92, E191, H268, D273,) in three distinct motifs (25). Active site residues are conserved among HCoV (**Fig 1A**). Our prior work had demonstrated that alanine substitutions of motif-I catalytic residues (D90A, E92A) were viable in MHV and SARS-CoV but had diminished proofreading and fitness, yet other groups failed to recover similar mutants with MERS-CoV and SARS-CoV-2 (12, 13, 25). To maximize the potential for viable virus recovery and to comprehensively query the genetic plasticity of the SARS-CoV-2 nsp14-ExoN active site, we performed saturation mutagenesis of motif I active site residues, generating all possible single through quadruple amino acid coding combinations at residues D90, E92, G93, and H95 (WT-DEGH) (**Fig. 1A**). Since residue 94 is cysteine in SARS-CoV and SARS-CoV-2, but alanine in other HCoV, the saturation mutagenesis libraries were generated holding this residue constant as cysteine or alanine totaling 320,000 potential combinations (**Fig. 1A**). Pooled library genomes were electroporated into BHK cells and co-cultured with Vero E6 cells overexpressing SARS-CoV-2 entry factors human transmembrane serine protease 2 (TMPRSS2) and angiotensin-converting enzyme 2 (ACE2), hereafter referred to as “VTA” cells. Cells were incubated at 33°C, and clarified viral supernatants were passaged on naive VTA cells to enrich for viable viruses. After passage, viral population sequences were obtained by the sequencing of viral RNA from infected cell monolayers, which revealed a constellation of possible mutational patterns in infectious viral genomes (**Fig. 1B**). A total of 124 unique variants were detected (**Table S1**). Over half of the variants had D90 or E92 mutated although no single mutants in either position were detected. 47 variants were detected with both D90 and E92 mutated all of which were also mutated at H95. A few of the variants that had both D90 and E92 mutated had changes that preserved the areas negative charge, with one variant switching the D and E residues, and several others with a D or E at one of the other residues. In addition, several mutants contained positively charged (R, K, H) or aromatic (Y, F) residues. All viable variants retained WT amino acid position V91. Position 95H was the most tolerant of change with a wide variety of amino acid substitutions detected. WT (DVEGCH) variants were not detected although sequences very close to WT (e.g. DVEGCT, DVEGCG) were detected at a low level (**Table S1**). A subset of mutants that were enriched after passage and/or that had 4 or more changes in motif-I (90-95aa) were selected for rescue via directed mutagenesis in the SARS-CoV-2 MA10 reverse genetic system (**Fig. 1C**) (26). Robust recovery was achieved for four of the nine mutant viruses: RAYF, AVFS, VHVV, and YQAV (WT = DEGH) (**Fig. 1D**). Using these optimized virus recovery conditions, we recovered virus containing the classical D90A/E92A substitutions in motif-I (AAGH) and then compared growth kinetics of the mutant panel and WT SARS-CoV-2 MA10 in VTA cells. All ExoN mutant panel viruses had significantly reduced replication compared to WT virus with ∼ 2 log reduction in peak titers at 36hr post infection (hpi) (**Fig. 1E**). To determine if ExoN- viruses were attenuated in a more biologically relevant and IFN competent culture system, we infected human primary airway epithelial (HAE) cell cultures with select mutant and WT viruses and monitored virus growth over time (**Fig. 1F**). HAE cells model the cellular complexity and architecture of the human conducting airway and contain epithelial subsets targeted by SARS-CoV-2 in humans (27, 28). Unlike WT SARS-CoV-2 MA10 virus which had increasing virus production over time, AAGH and YQAV viruses were significantly attenuated in HAE. Altogether, our saturation mutagenesis approach coupled with technical experimental improvements in virus culture demonstrate that multiple constellations of ExoN Motif I mutations are viable in SARS-CoV-2, but resultant viruses are attenuated for replication in Vero cells and significantly attenuated in primary human airway epithelial cells.

**Figure 1:**
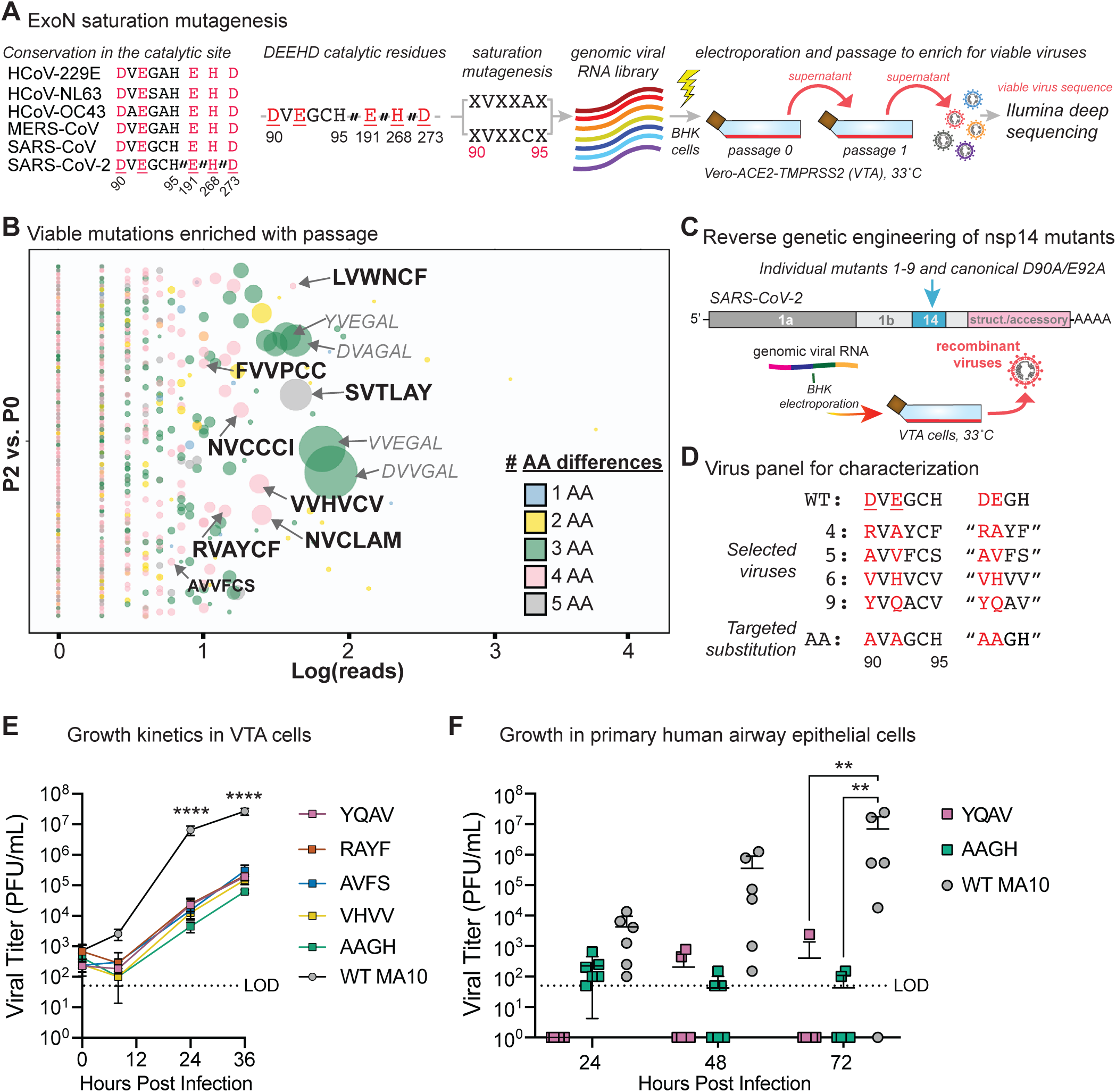
SARS-CoV-2 nsp14-ExoN mutants are viable but attenuated for replication in vitro. **(A)** Schematic of the saturation mutagenesis of ExoN motif-I. A viral RNA library with all possible combinations of amino acid mutations at positions 90, 92, 93 and 95 was electroporated into BHK cells and cocultured with VTA cells at 33°C. Viable viruses were enriched via passage on VTA cells and identified through Illumina deep sequencing. **(B)** Frequency of mutation series in viable genomes after passage. **(C)** Schematic for reverse genetic approach to engineer sets with 4 or more changes from WT as well as the canonical D90A/E92A mutation into SARS-CoV-2 MA10. **(D)** List of substitutions present within the ExoN motif of panel viruses. **(E)** Growth kinetics of SARS-CoV-2 MA10 ExoN mutant panel. VTA cells were infected at an MOI of 0.01, input removed, monolayers were rinsed once and infectious virus in culture media was measured via plaque assay over time. Asterisks indicate significant differences (P <0.0001) among mutant viruses and WT via Two-way ANOVA Dunnet’s multiple comparison test. **(F)** Growth of select SARS-CoV-2 MA10 ExoN viruses and WT in primary human airway epithelial (HAE) cell cultures. Data is combined from two independent studies with cells derived from two different human donors. HAE were infected at an MOI of 0.1 for 1.5hr after which input was removed and cultures were washed with PBS. Infectious virus production was measured via plaque assay of apical washes collected at the indicated times. Asterisks indicate statistical differences titer via Mann-Whitney test (YQAV P = 0.0054, AAGH P = 0.0076).

To gain insight into the mechanism of replication attenuation, we performed a series of biochemical and virologic studies. First, we assessed the exonuclease activity of select ExoN mutants in a FRET-based biochemical assay with nsp10-14 fusion protein, since nsp10 is a required co-factor of ExoN (29, 30). For this assay, the nsp-10-14 enzyme attacks a double stranded RNA substrate with mismatched base pairing liberating the fluorophore and quencher associated with the probe resulting in measurable fluorescence. While WT nsp14/nsp10 exhibited notable exonuclease activity, none of the mutant enzymes assessed, including the canonical AAGH, had any detectable exonuclease activity (**Fig. S1)**. Thus, substitutions observed in viable SARS-CoV-2 MA10 viruses rendered the nsp14-ExoN enzyme catalytically dead.

### SARS-CoV-2 ExoN mutations decrease specific infectivity

Next, we sought to determine the impact of ExoN inactivation on SARS-CoV-2 replication, RNA synthesis and specific infectivity. For these studies, we generated the canonical motif-I AAGH mutation in the SARS-CoV-2 WA/1 background given the abundance of data we have previously generated with this approach with MHV and SARS-CoV (12, 13, 31). First, we recovered WT and AAGH viruses at either 33° in VTA cells (P01) and then generated P1 stocks by passaging on VTA cells at either 33°C or 37°C. We then evaluated viral growth kinetics of these stocks at the temperature in which the stocks were prepared normalizing virus input by infectious titers determined on VTA cells (MOI = 0.01). While recovery of ExoN- viruses was only achieved 33°C, the levels and kinetics of replication were similar at both 33°C and 37°C for WT and AAGH viruses (**Fig. 2A**). Similar to what we had demonstrated with SARS-CoV-2 MA10 ExoN- viruses above, SARS-CoV-2 WA/1 AAGH virus was attenuated for replication in VTA cells with significant differences from WT 24-48hpi. Mean infectious titer differences between WT and AAGH viruses were greatest at 24hpi, exceeding 3 logs for both temperature conditions. We then performed RT-qPCR to quantitate viral genomic RNA in this growth kinetic to better understand the relationship between infectious virus and virus particle production. In contrast to infectious titers, viral RNA copy number was similar for WT and AAGH viruses at early times but diverged beginning at 24hpi which continued through 48hpi (**Fig. 2B**). Like infectious titer data, temperature did not have an appreciable difference on the numbers of secreted viral RNA genomes for WT and AAGH viruses (**Fig. 2B**). Using the viral RNA copy number (**Fig. 2B**) and infectious titer (**Fig. 2A**) data, we then calculated the specific infectivity (i.e. PFU/ Genome RNA copy ratio), a value which provides insight into the relative infectiousness of a virus sample.

**Figure 2:**
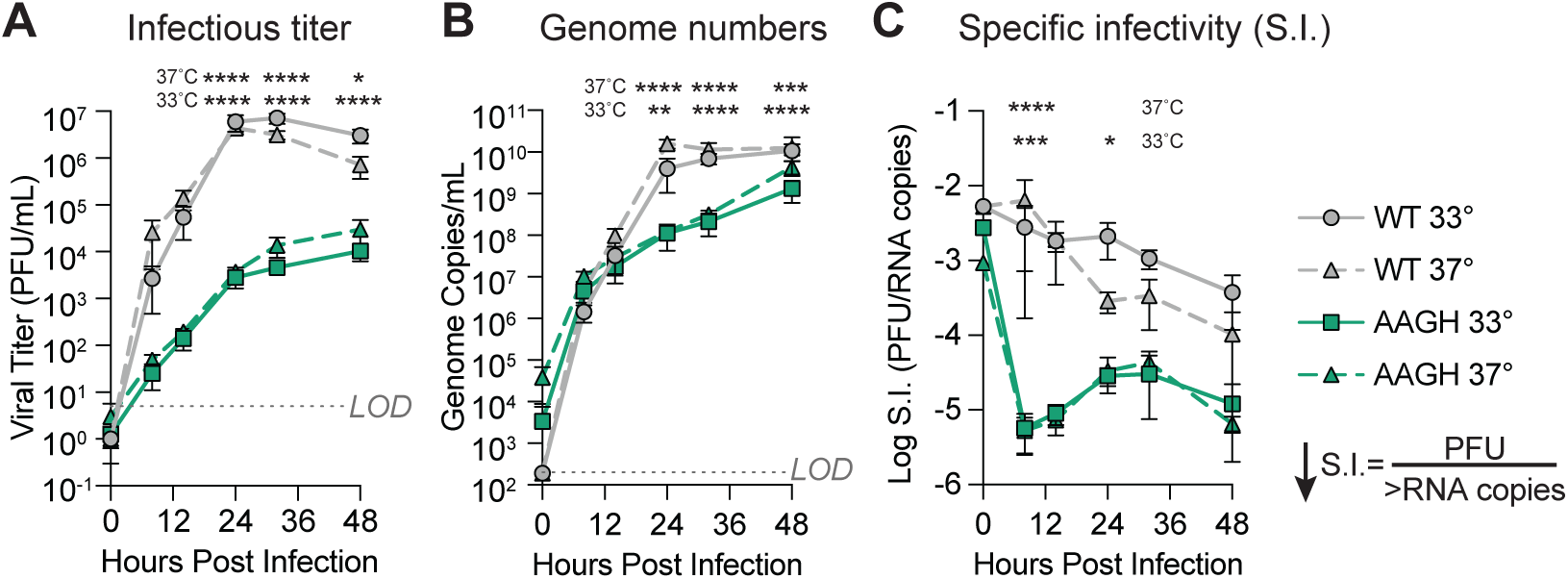
SARS-CoV-2 ExoN inactivating mutations abolish enzymatic activity and decrease specific infectivity. **(A)** The *in vitro* exonuclease activity for select ExoN mutants was determined by a fluorescence RNA cleavage assay using nsp10/nsp14 fusion proteins. The RNA cleavage activity of WT and mutant nsp14 fusion proteins was measured over time using normalized amounts of protein and double stranded RNA FRET robe. Symbols represent mean values and ± SEM error bars from four independent experiments. **(B)** Growth of SARS-CoV-2 WT or AAGH virus at 33° or 37°C in VTA cells infected at an MOI of 0.01 PFU/cell. Titers were determined in supernatant by plaque assay at the indicated times. **(C)** Viral genome copy number in samples evaluated in “B” was determined by RT-qPCR. **(D)** Specific infectivity was determined as the ratio of genome copies/PFU from data shown in “B” and “C”. Graphed are mean values ± standard error of the mean (SEM) from triplicate infections of two independent experiments. Asterisks indicate statistical significance by Two-Way ANOVA with Sidak’s multiple comparison test.

While the specific infectivity (S.I.) varied over time for both WT and AAGH virus samples, generally AAGH had a 10-100-fold decreased S.I. consistent with AAGH generating more non-infectious RNA or particles relative to WT virus (**Fig. 2C**). Overall, these studies show that decreased temperature was critical for the recovery of SARS-CoV-2 ExoN-, but temperature does not have a significant impact on WT or ExoN- viral replication once stocks are established. These data demonstrate that mutations inactivating SARS-CoV-2 nsp14-ExoN attenuate replication, RNA synthesis, and infectious virus production and that in the absence of a functional nsp14-ExoN, the numbers of non-infectious particles is decreased.

### SARS-CoV-2 AAGH has diminished competitive fitness

We next performed head to head competition studies with SARS-CoV-2 WT and AAGH ExoN- mutant viruses to assess competitive fitness in low MOI infections (0.01) (**Fig. 3A**). Competition studies directly compare replicative fitness of two or more viral variants to accurately benchmark comparative replicative fitness. We mixed WT or AAGH virus at infectious particle ratios of 1:1 or 1:9, the latter condition providing AAGH virus in vast excess to WT. We infected VTA cells with these mixtures (passage 1, P1) at either 33° or 37°C and allowed the infection to progress for 18hr after which these P1 viral supernatants were blindly passaged to naive VTA cells for another 18hrs. Competition at both temperatures was evaluated to determine if temperature impacted competitive fitness. The ratio of WT and AAGH genomes present at the end of P1 and P2 was determined by RT-PCR followed by area under the curve analysis of Sanger sequencing trace results at each of the engineered codons in nsp14 (**Fig. 3B**). ExoN- AAGH viruses failed to compete with WT viruses when mixed at a 1:1 ratio with the majority of the population consisting of WT sequences after a single passage. Even when introducing significantly more AAGH virus at the start of passaging (1:9, WT: AAGH), AAGH virus comprised less than 50% of the population after P1 and by the end of P2, was less than 5% of the population. While the AAGH virus appeared to have increased fitness at 33°C relative to 37°C after 1 passage when mixed at a ratio of 1:9 (WT:AAGH), the AAGH virus was still significantly outcompeted by WT regardless of temperature. These results indicate that even when given a significant advantage in terms of input MOI or temperature, SARS-CoV-2 ExoN- virus is unable to compete with WT virus.

**Figure 3:**
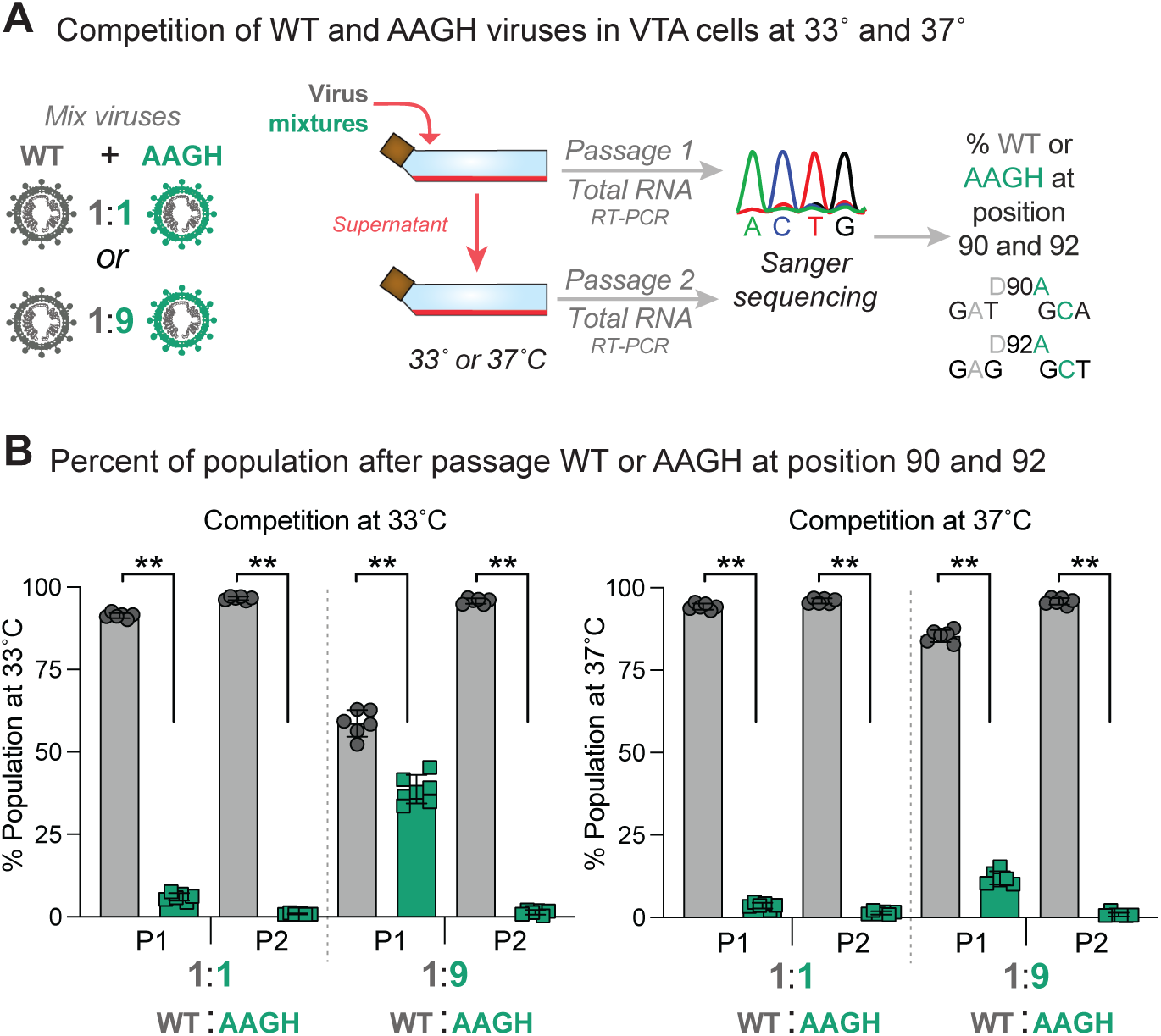
SARS-CoV-2 AAGH virus has diminished competitive fitness. **(A)** Competitive fitness study schematic. VTA cells in 6-well plates were co-infected with three independent lineages of SARS-CoV-2 WT mixed with SARS-CoV-2 AAGH at 1:1 MOI ratios of 0.005 PFU/cell or 1:9 MOI ratios of 0.001 (WT) and 0.009 (AA) PFU/cell for 1hr, input virus was removed, monolayers were washed twice and growth medium was added. After 18hr, supernatants (Passage 1, P1) were blindly passaged on VTA cells and total RNA was collected in TRizol. After 18hr, total RNA was similarly collected from P2 monolayers. RT-PCR amplicons encompassing nsp14-ExoN motif I were analyzed by Sanger sequencing. **(B)** Percent population WT or AAGH after passage. Sanger sequencing traces were analyzed for area under the curve at nucleotide positions 18,308 (D90A; WT G<u>A</u>T, AAGH G<u>C</u>A) and 18,314 (E92A; WT G<u>A</u>G, AAGH G<u>C</u>T) to determine the percent of the population of WT or ExoN- genotype. The error bars represent the mean and standard deviation. The asterisks indicate statistical significance by Mann-Whitney test.

### SARS-CoV-2 AAGH virus has increased sensitivity to nucleoside analogues

We have previously demonstrated that both MHV and SARS-CoV nsp14 exonuclease mutants are more sensitive to antiviral and mutagenic nucleoside analogues compared to WT viruses, functionally demonstrating the consequences of proofreading defects on replication (14, 32, 33). To determine if SARS-CoV-2 ExoN- viruses are similarly sensitive to nucleoside analogues, we performed antiviral drug assays with SARS-CoV-2 WT and AAGH nanoluciferase reporter viruses in VTA cells (**Fig. 4**). We compared β-d-N4-Hydroxycytidine (NHC, EIDD-1931) the parental nucleoside to the prodrug molnupiravir; GS-441524, the parental nucleoside of the prodrug remdesivir; and 5-fluorouracil (5-FU), a known mutagenic small molecule nucleoside analog. As we had observed for MHV and SARS-CoV ExoN- viruses, SARS-CoV-2 AAGH had increased sensitivity to all small molecules assessed (14, 32, 33). Unlike WT virus which was refractory to the effects of 5-FU even at concentrations up to 400µM, SARS-CoV-2 AAGH was sensitive to treatment (EC_50_ = 124.7 µM). Similarly, SARS-CoV-2 AAGH had increased sensitivity to both NHC and GS-441524 with 6 and 10-fold shifts in antiviral potency, respectively, suggesting an intrinsic role in defense against antiviral nucleoside analogues (34). Thus, without a functional ExoN, SARS-CoV-2 has a notable increase in sensitivity to antiviral agents and mutagenic nucleoside analogues demonstrating the impact of ExoN mediated proofreading on nucleoside analogue sensitivity.

**Figure 4:**
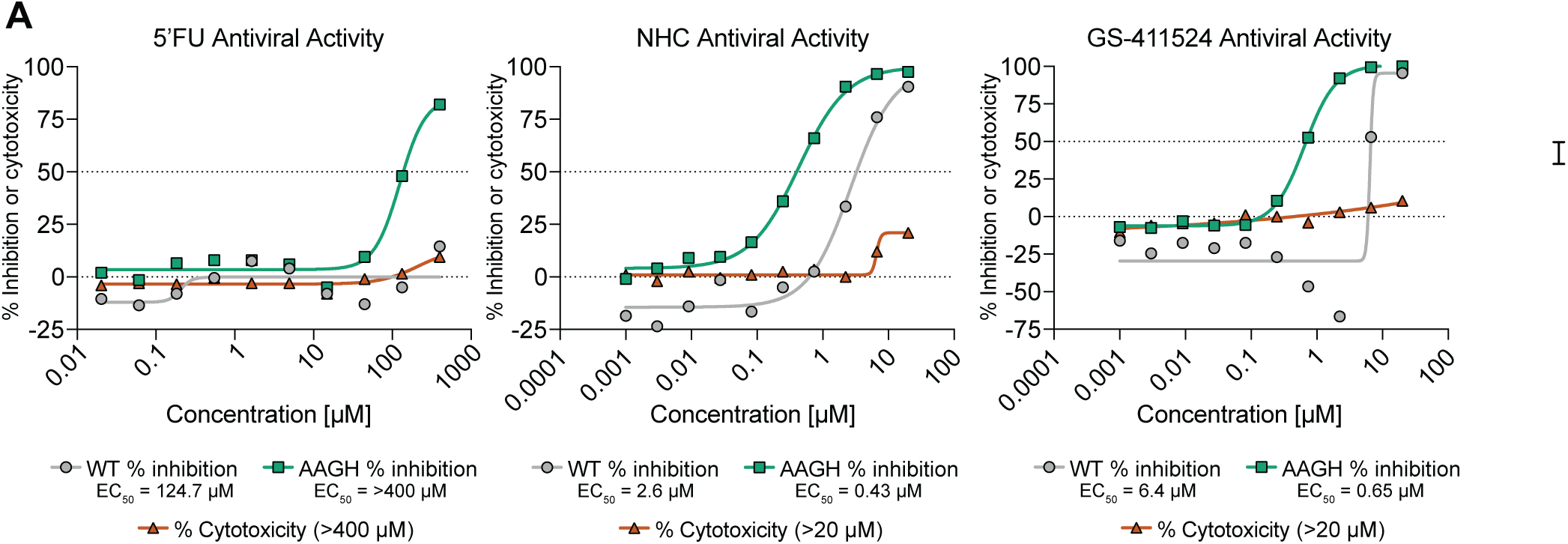
SARS-CoV-2 AAGH virus has increased sensitivity to nucleoside analogues. VTA cells were infected with nanoluciferase-expressing SARS-CoV-2 WT or AA at 37°C at an MOI of 0.1 for 1hr after which input was removed, monolayers were washed and then exposed to a dose response of 5-fluorouracil (5-FU), β-d-N4-Hydroxycytidine (NHC, EIDD-1931) or GS-441524 in Infection Media. Concurrently, non-infected cells were treated similarly to determine cytotoxicity. After 24hr, viral replication was assessed by NanoGlo Luciferase Assay System (Promega) and cytotoxicity was determined by CellTiterGlo Assay (Promega). Each condition was evaluated in triplicate in two independent studies. Values were normalized to the uninfected and infected vehicle DMSO controls (0 and 100% infection, respectively). Data were fit using a four-parameter nonlinear regression analysis using GraphPad Prism. EC50 and CC50 (cytotoxic concentration at which 50% of cells are viable) values were then determined as the concentration reducing the signal by 50%.

### SARS-CoV-2 ExoN- viruses have decreased replication fidelity

The loss of nsp-14 ExoN proofreading activity can manifest in an accumulation of mutations measurable by deep sequencing (12, 13, 23). Thus, we performed similar studies with select SARS-CoV-2 ExoN- loss of function viruses. First, we infected VTA cells with WT or SARS-CoV-2 AAGH virus at a low multiplicity of infection and harvested total RNA for Illumina RNAseq analysis when the majority of the monolayer exhibited a cytopathic effect (CPE). Sequencing read depths were similar for WT and AAGH viruses at the 5’ and 3’ ends of the genome but appeared to drop slightly for AAGH in the middle of ORF1a through to the M gene (**Fig. 5 A-B**). Nevertheless, 99.9% of the AAGH genome had a sequencing coverage greater than 1000X. An increased number of variants were detected in AAGH virus samples as compared to WT virus (**Fig. 5A-B**). Congruent with slight differences in read depth noted above, AAGH samples for sequencing had fewer copies of viral genomes measured by RT-qPCR than WT viral samples (**Fig 5C**). In agreement with data above, the mutation frequency of AAGH virus was significantly increased compared to WT virus (**Fig. 5D**) with notable increases in both transition and transversion mutations (**Fig. 5E-F**). While we had shown above that SARS-CoV-2 AAGH virus had increased susceptibility to mutagenic nucleoside 5-FU, we next sought to determine if growth of ExoN- virus in the presence of 5-FU would result in an accumulation of variant mutations as we had previously shown with SARS-CoV(14). We infected VTA cells with WT or SARS-CoV-2 AAGH or YQAV viruses in the presence of DMSO vehicle or 100µM 5-FU. After 24hr, viral RNA was isolated from clarified supernatants for deep sequencing analysis. SARS-CoV-2 ExoN- viruses grown in the presence of a mutagenic small molecule 5-FU accumulate mutations at a greater rate (**Fig. 5G**) than the same viruses grown in media containing vehicle, thus demonstrating the functional consequence of diminished replication fidelity and proofreading activity. Lastly, consistent with our prior report that MHV nsp-14 ExoN mediates viral RNA recombination (35), similar analysis of our deep sequencing data indicated that SARS-CoV-2 AAGH had altered viral RNA recombination patterns (**Fig. S2)**. Altogether, we demonstrate that SARS-CoV-2 nsp14-ExoN is key to replication fidelity, proofreading and RNA recombination.

**Figure 5:**
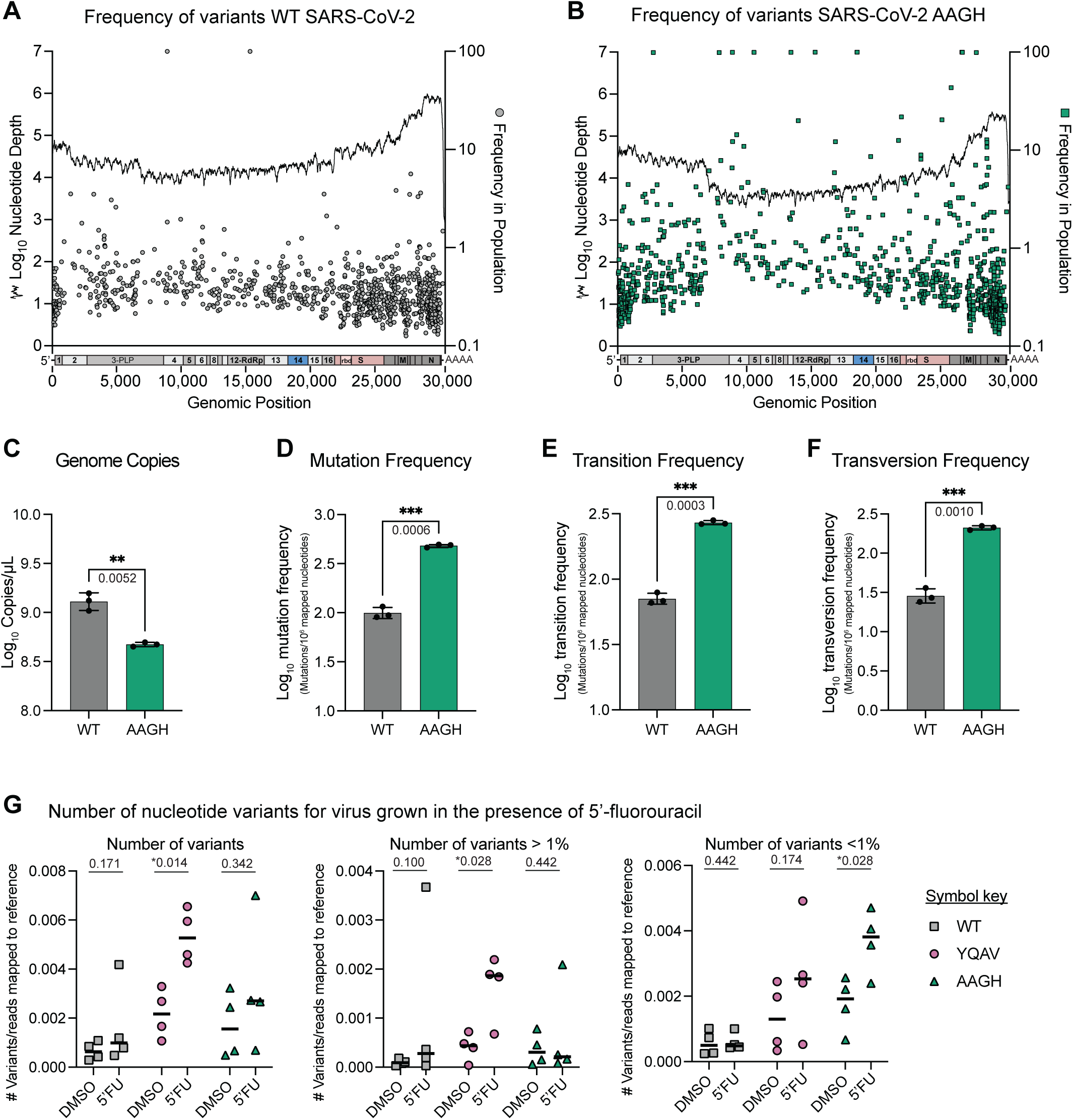
SARS-CoV-2 ExoN- viruses have decreased replication fidelity. **(A-B)** Nucleotide variant identification by deep sequencing. VTA cells were infected with SARS-CoV-2 WT or AAGH at an MOI of 0.01 in biological triplicate at 33° until 70% of the monolayer exhibited cytopathic effect. Total RNA was isolated and variants were identified by RNAseq. Sequence coverage, variant frequency and location on the viral genome is shown for representative samples of WT (A) and AAGH (B) viruses. **(C)** Viral genome concentration in total RNA analyzed in “A” and “B” by RT-qPCR. **(D)** Total mutation frequency. **(E)** Transitions frequency. **(F)** Transversion frequency. For **C**, viral genome copies were log transformed. For **D-F**, the ratio of mutations per 1 million mapped nucleotides was generated and log transformed. For **C-F**, the symbols represent biological replicates, line represents the mean, error bars signify the standard deviation, and statistical significance was determined by one-tailed Welch’s t-test. Asterisks indicate statistical differences and p-values are displayed. **(G)** Mutation frequency of WT and ExoN- viruses in the presence of 5’-fluorouricil. VTA cells were infected with SARS-CoV-2 WT, AAGH or YQAV in the presence of DMSO vehicle or 100µM 5’-FU at 33°for 24hr after which clarified supernatants were harvested, RNA extracted and analyzed by Illumina MiSeq. Each condition was performed in quadruplicate. The total number of variants, the number of variants > 1% and <1% normalized to the number of reads mapped to the reference sequence is shown. Statistical significance determined by Mann-Whitney test is denoted by an asterisk and p-values are displayed.

### SARS-CoV-2 ExoN- viruses are attenuated in vivo

We next evaluated the replicative fitness and pathogenic potential of SARS-CoV-2 ExoN- viruses in animal models. Since we generated ExoN mutant viruses in the mouse adapted MA10 genetic background, we first evaluated pathogenesis in 10-week old BALB/c mice infected with 6E+05 PFU SARS-CoV-2 MA10 WT or SARS-CoV-2 MA10 YQAV virus. The YQAV mutant was chosen for these studies because the stock titers for other mutants were not high enough to match the 6E+05 of WT. Unlike WT virus which caused progressive body weight loss over time, SARS-CoV-2 MA10 YQAV infection did not cause weight loss during this study (**Fig. 6A**). Concordant the inability to cause body weight loss, SARS-CoV-2 MA10 YQAV replication in lung tissue was significantly decreased by three to four logs, as compared to WT virus titers on 1, 2 and 4 days post infection (dpi) (**Fig. 6B**). Since SARS-CoV-2 MA10 YQAV was not pathogenic in BALB/c mice, we then sought to evaluate ExoN- virus pathogenesis in a highly susceptible mouse model, C57BL/6 mice with hACE2 expressed via the epithelial K18 promoter (“K18” mice). In addition, we sought to evaluate our entire panel of ExoN- SARS-CoV-2 to more comprehensively assess pathogenic potential. We infected 8-9 month old male and female K18 mice with ∼3E+04 PFU of SARS-CoV-2 MA10 ExoN- panel viruses. Similar to infection in BALB/c, WT virus infection caused progressive weight loss in K18 mice yet ExoN- viruses infection failed to induce body weight loss (**Fig. 6C**). Gross lung pathology was evident in only WT infected mice on 6dpi (**Fig. 6D**). ExoN- virus replication in the lung at both 2 and 6dpi was significantly attenuated as compared to WT virus in K18 mice with mean titers differing by approximately 4 logs (**Fig. 6E, F**). Of ExoN- viruses, only RAYF had measurable virus in the lung on 6dpi suggesting there may be subtle differences in replicative fitness *in vivo* among the panel viruses (**Fig. 6F**). Lastly, mean brain titers differed by approximately 9 logs with evident neuroinvasion and replication in most WT infected animals while the brain titers of all ExoN- viruses were below the limit of detection (**Fig. 6G**). Altogether, these data indicate that SARS-CoV-2 MA10 ExoN- viruses are severely attenuated for replication and pathogenesis in WT mice and in the highly susceptible K18 hACE2 mouse model.

**Figure 6:**
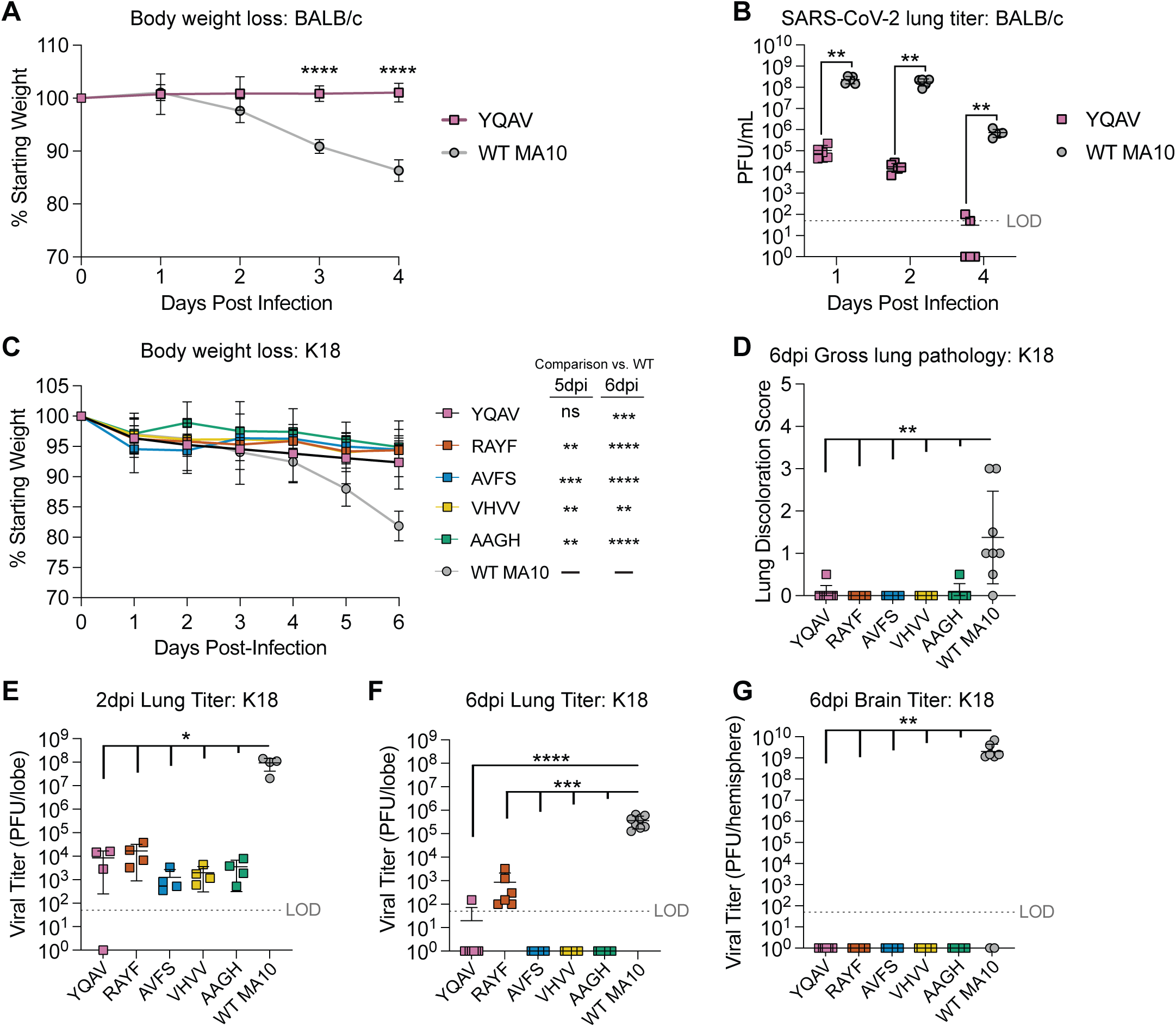
SARS-CoV-2 ExoN- viruses are attenuated in multiple mouse models. **(A)** Body weight loss in BALB/c mice. 10-week-old female BALB/c mice were intranasally infected with 6.8E+05 PFU of WT SARS-CoV-2 MA10 (N = 15) or related YQAV virus (N = 15) and body weight was measured daily. **(B)** Viral lung titer in BALB/c mice. On the indicated day, the bottom right lung lobe was harvested from animals described in “A” and SARS-CoV-2 titers were determined on VTA cells by plaque assay. LOD = Limit of detection. **(C)** Body weight loss in C67BL/6 K18 hACE2 mice. 8-9-month-old male and female C57BL/6 K18-hACE2 mice were infected with 3.0E+04 PFU of WT SARS-CoV-2 MA10 (N = 12) or related ExoN- viruses AAGH (N = 12), RAYF (N = 10), AVFS (N = 10), VHVV (N = 10), or YQAV (N = 12) after which body weight was measured daily. **(D)** Gross lung pathology. SARS-CoV-2 infection can cause a hemorrhage-like lung discoloration scored on a scale of 0 (normal) to 4 (100% discolored). **(E-G)** Viral titers in the lung on 2dpi **(E)** and 6dpi **(F)** or brain on 6dpi **(G)**. For the longitudinal body weight data, statistical significance was determined using a Mixed Effects Analysis Model with Multiple Comparisons. For the lung hemorrhage score, and lung and brain titer data, statistical significance was determined via a one-tailed, Mann-Whitney test. Asterisks indicate statistical differences (ns=not significant, * p value<0.05, ** p value<0.01, *** p value<0.001, **** p value<0.0001).

### Interferon deficiency partially restores SARS-CoV-2 ExoN- replicative fitness in vivo

To determine if IFN signaling was playing a role in the attenuation of SARS-CoV-2 Exon- replication in mice, we first assessed SARS-CoV-2 ExoN- virus susceptibility to IFN pretreatment in vitro (**Fig. 7A**) (36). The potency of IFN-ß in VTA cells was nearly 20-fold greater against SARS-CoV-2 AAGH virus (IC_50_ = 0.019 ng/mL) compared to WT virus (IC_50_ = 0.34 ng/mL) thus confirming the increased IFN sensitivity of SARS-CoV-2 ExoN- viruses. We found that SARS-CoV engineered to have the exact same ExoN-inactivating AAGH mutation also exhibited increased sensitivity to IFN-ß (**Fig. S3**) (23).

**Figure 7:**
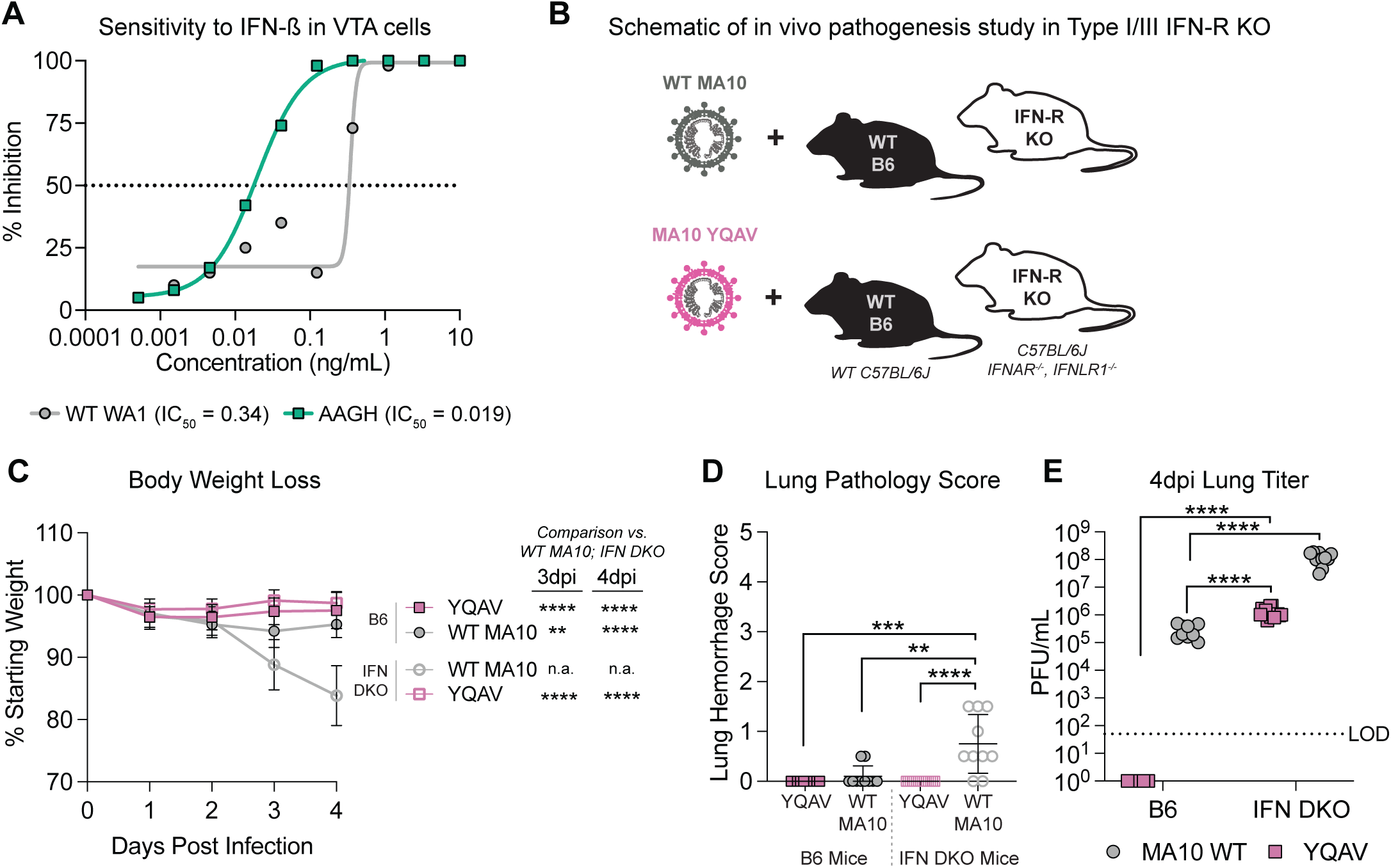
Interferon deficiency partially restores SARS-CoV-2 ExoN- replicative fitness in vivo. **(A)** Sensitivity to interferon beta in VTA cells. VTA cells were pre-treated with dose response of IFNβ for 24hr. Interferon was removed and then cells were infected with SARS-CoV-2 WT or AAGH nanoluciferase-expressing viruses an MOI of 0.1. The plates were incubated at 37°C for 1hr, input virus was removed, cultures were washed and growth media was added. After 72hr at 37°C, viral replication was mediated by NanoGlo Assay. **(B)** Schematic of type I and III Receptor KO mouse experiment. 8-15 week old male and female WT or interferon αβλ receptor deficient mice were infected with 6.8E+05 PFU of either WT SARS-CoV-2 MA10 (WT N = 10, KO N = 11) or YQAV virus (WT N = 10, KO = 15). **(C)** Body weight loss over time of mice described in “B”. **(C)** Gross lung pathology score for mice described in “B”. **(D)** Viral lung titer 4dpi via plaque assay. LOD = limit of detection. Asterisks indicate statistical significance by Two-way ANOVA with a Dunnet’s multiple comparison test in “C” and a Two-way ANOVA with a Tukey’s multiple comparison test in “C” and “D”.

To elucidate the importance of IFN signaling in replicative fitness and pathogenic potential *in vivo*, we infected WT mice or those deficient in type I (alpha/beta, α/β) and type III (lambda, λ) IFN receptor (Type I/III IFN-R KO) with either WT SARS-CoV-2 MA10 or SARS-CoV-2 MA10 YQAV virus (**Fig. 7B**). Like the above studies, WT infection caused progressive body weight loss, yet this metric was not affected by ExoN- virus infection (**Fig. 7C**). Only WT infection caused measurable gross lung pathology on 4dpi (**Fig. 7D**). While SARS-CoV-2 MA10 YQAV infectious titers in WT mice were below the limit of detection (LOD) on 4dpi, remarkably, they were approximately 4.5 logs above the LOD in type I/III IFN-R KO mice and exceeded that achieved by WT virus in WT mice at similar times (**Fig. 7E**). WT viral replication increased in type I/III IFN-R KO mice as well, although only by approximately 3 logs (**Fig. 7E**). We next performed similar studies with SARS-CoV MA15 WT and ExoN-virus (i.e. AAGH) in WT mice and those deficient in Type I and Type II IFN signaling (**Fig. S3**). As expected, WT SARS-CoV MA15 virus infection in either strain of mouse caused progressive body weight loss with mice meeting humane endpoints for euthanasia by 4dpi (**Fig. S3**). Although weight loss and clinical disease was significantly attenuated in SARS-CoV MA15 AAGH infected WT mice, increased weight loss was observed in type I/II IFN-R deficient mice indicating a partial restoration of pathogenesis. Early in infection, SARS-CoV MA15 AAGH viral replication was decreased ∼2 log in the lungs either mouse strain compared to WT virus (**Fig. S3**). At 4dpi, viral replication for both WT and SARS-CoV MA15 AAGH was increased ∼2 log in type I/II IFN-R deficient mice compared to WT. Altogether, these studies indicate that SARS-CoV-2 ExoN- viruses do not have a generalizable defect in replication that limits replicative capacity *in vivo*. In addition, these data demonstrate that IFN signaling plays a significant role in attenuating replication and pathogenesis of both SARS-CoV and SARS-CoV-2 ExoN- suggesting that ExoN function in virus and host innate immune interactions are important for replicative fitness and pathogenesis in vivo.

## Discussion

RNA viral replication infidelity and capacity for genetic change is essential for the rapid adaptation to new hosts, evasion of immunity and to develop therapeutic resistance. CoV replication and transcription, although largely driven by the RdRp, is ultimately a coordinated effort among RdRp and a variety of viral non-structural proteins that make up the RTC. While other RNA viruses, like Lassa virus, encode proteins with exonuclease activity (37), CoV are unique in that their viral exonuclease (ExoN, nsp14) exhibits proofreading function which is thought to enhance replication fidelity and facilitate the maintenance of their relatively large RNA genomes (9, 12–14). Although CoV ExoN-mediated proofreading was first demonstrated via reverse genetic ablation and in vitro biochemical studies on SARS-CoV and MHV ExoN activity previously(10, 12, 13), attempts to generate similar ExoN- mutants with *Alphacoronavirus*, TGEV and HCoV-229E, or Beta-CoV, MERS-CoV and SARS-CoV-2, have been reported to fail (10, 25, 38). While demonstrating a critical role of ExoN in CoV replication and viability, the mechanisms influencing dramatically different experimental outcomes suggest potential differences in ExoN function across the CoV family. Coupling reverse genetics, saturation mutagenesis and specific culture conditions, we generated a large library of viable ExoN Motif I mutant viral RNAs, thus revealing significant amino acid heterogeneity is tolerated within the ExoN motif I of viable viruses and that certain residues are less tolerant of change (V91) than others (H95). In addition, we optimized the cellular environment for recombinant viral recovery. First, we utilized the highly transfectable baby hamster kidney cells (BHK) for electroporation and then cocultured the transfected BHK with a highly permissive Vero E6 cell overexpressing key viral entry factors, ACE2 and TMPRSS2 (i.e. VTA cells). Unlike SARS-CoV-2 cytopathic effect in typical Vero cells, SARS-CoV-2 replication in VTA cells causes syncytia which in the context of recombinant viral recovery may promote cell to cell viral spread. In addition, both BHK and Vero E6 have known defects in Type I IFN signaling (39, 40). Second, we had preliminary data indicating that recovery of debilitated viruses could be improved at lower temperature (33°C) surmising that lower temperature may slow cell division, alter viral protein structure/function and/or diminish innate immune function the latter which has been shown in the context of Chikungunya virus and rhinovirus infection (41, 42).

Collectively, our saturation mutagenesis approach coupled with multiple changes in the cellular environment facilitated the recovery of multiple SARS-CoV-2 viruses lacking a functional ExoN revealing a remarkable genetic pliability of the SARS-CoV-2 ExoN active site pocket. These advances also facilitated the recovery of virus with the canonical D90A/E92A (AAGH) mutation. Interestingly, we found that once recovered, SARS-CoV-2 ExoN- viruses grew similarly at 33° and 37°C in VTA cells indicating they did not have a temperature sensitive growth phenotype and that low temperature was more likely creating more hospitable cellular milieu that potentiates recovery. These new approaches may facilitate the rescue of other previously non-recoverable, LOF mutants of SARS-CoV-2 and other CoVs, expanding the viral genetic tools available for the study of CoV replication and pathogenesis.

The evolution and retention of the CoV nsp14-ExoN suggest it provides a positive functional benefit for survival, driven by improved fidelity, protection from natural cellular antiviral nucleosides perhaps like 3′-deoxy-3′,4′-didehydro-cytidine 5′-monophosphate (ddhCMP) as well as other less well defined innate immune antagonistic functions (34, 43–45). The generation of LOF ExoN deficient CoVs enables the investigation of ExoN function in replication and pathogenesis and provides the basis for a pan-CoV live attenuated reversion resistant vaccine strategy (23, 46). Although SARS-CoV and SARS-CoV-2 have genomes that are approximately 80% identical, their nsp14 proteins are 95% identical (47). Despite these genetic similarities, there are both similarities and differences among SARS-CoV and SARS-CoV-2 ExoN-virus phenotypes. Aside from contrasting technical approaches for recombinant virus recovery, SARS-CoV-2 ExoN- is more attenuated for replication in Vero cells than SARS-CoV ExoN whose replication mirrors that of WT virus for the first 24hr of infection (23). Although performed in different cells, MHV ExoN- virus growth in murine DBT cells is significantly attenuated more akin to SARS-CoV-2 ExoN- described herein (12). Differences in replicative fitness aside, ExoN deficient SARS-CoV, SARS-CoV-2 and MHV all exhibit a loss of replication fidelity exhibited by the accumulation of mutations in viral RNA during replication (12–14, 23). Relatedly, the specific infectivity of viral particles is diminished in SARS-CoV-2 and MHV ExoN- viruses driven by the production of increased levels of defective RNA and non-infectious particles (14). Increased susceptibility to both nucleoside antiviral drugs like remdesivir and mutagenic nucleosides like 5-FU demonstrate shared functional defects in proofreading activity amongst MHV, SARS-CoV and SARS-CoV-2 ExoN- viruses (14, 32). These data highlight that despite some ability of nsp14-ExoN to excise non-native nucleosides, antivirals like remdesivir overwhelm the proofreading activities of the RTC to exert their antiviral effect. CoV subgenomic transcription is mediated by a highly regulated RNA recombination mechanism called discontinuous transcription (48). While viral RNA recombination is required during CoV replication, it has also been hypothesized to facilitate host range expansion through the exchange of spike gene sequences (49, 50). Recent reports implicate nsp14-ExoN in viral RNA recombination during replication and transcription, although the governing molecular mechanisms remain unclear. Like MHV ExoN- virus, SARS-CoV-2 ExoN- viruses have altered global recombination patterns providing additional evidence that CoV exonuclease plays a role in viral RNA recombination (35). Lastly, we recovered multiple viable viruses with a variety of substitutions at nsp14 residues D90, E92, G93, and H95 where D90 and E92 are motif I active site residues and G93 and H95 interact with 3’ end nucleotides in the product strand (22). Simultaneous changes to all four amino acid residues noted above thus represent potentially significant structural and mechanistic alterations to nsp14-ExoN activity. The fact that single mutants at either D90 or E92 were not detected after passage and that the alteration of these residues was associated with additional amino acid changes suggest that this region may have functions beyond canonical exonuclease activity. It is especially interesting that charged residues were enriched in motif I of ExoN when absent in the typical D90/E92 positions. Given that nsp14 plays a central role in the multi-protein RTC complex, and engages nsp10 and 16, aside from inactivating nsp14-ExoN function, these changes may impact other important RTC functions (11, 51). In addition, recent biochemical data indicate that nsp8, an essential cofactor of the RdRp-nsp7-nsp8 complex, enhances nsp14-10 exonuclease activity providing insight into the complexity of RTC protein interactions and functions (52). Thus, the attenuation of SARS-CoV-2 ExoN- replication may in part be driven by a combination of ExoN intrinsic effects on replication fidelity and extrinsic effects on viral RTC protein interactions and functions.

The majority of the literature on CoV nsp14 and the host innate immune response relates to its N7-Mtase activity which plays an essential role with nsp16 2’-O-methyltransferase in generating a 5’ RNA cap structure that is indistinguishable from that on host mRNAs (53, 54). Without these modifications, viral genomic and subgenomic RNA would be ripe targets for RNA pattern recognition molecules like MDA5 and the induction host antiviral programs like IFN (55). Remarkably, only a handful of studies have reported a potential link between nsp14 exonuclease function and the innate immune response. MHV ExoN- is attenuated for replication and has increased sensitivity to IFN pretreatment in a murine astrocytoma cell line (i.e. DBT) (36). Interestingly, with serial passage of MHV ExoN-, compensatory mutations that restore replicative fitness are gained yet ExoN enzyme remains inactive and increased IFN sensitivity was preserved (36, 46). These studies suggest a function for ExoN 3’-5’ exonuclease activity in IFN antagonism. A recent overexpression study demonstrated that both ExoN and N7-Mtase domains of SARS-CoV-2 nsp14 diminish IFN stimulated gene (ISG) translation suggesting that nsp14 may have direct host modulatory activities (44). In addition, mutation of nsp14-ExoN zinc fingers (ZF) for the *Alphacoronavirus* TGEV resulted in WT levels of replication but reduced apoptosis and IFN-ß production further implicating nsp14 in modulation of the host response(38). Genetically unrelated Lassa virus NP protein has exonuclease activity which digests viral double stranded RNA, thereby diminishing the sensing of this molecular pattern instigator of RIG-I, an RNA sensing inducer of the innate antiviral response (37, 56). If CoV ExoN similarly antagonize the host response through the digestion of viral RNA PAMPs, this function would not be detected in overexpression studies in the absence of viral RNA synthesis. Nevertheless, there is a growing body of literature indicating that viral exonucleases can exert key functions in replication while also serving to antagonize the innate immune response. Future studies are aimed at addressing the underlying molecular mechanisms governing host innate immune antagonism of Sarbecovirus ExoN.

Here, we show that SARS-CoV and SARS-CoV-2 ExoN- viruses are more sensitive to IFN pretreatment *in vitro* and are significantly attenuated in WT mice but attenuation *in vivo* can partially be restored via knock out of IFN signaling. SARS-CoV-2 ExoN- virus replicates robustly in Type I/III IFN-R KO mice achieving titers at 4dpi like that of SARS-CoV-2 WT virus in WT mice at the same timepoint. Given that we utilized either Type I/III IFN-R KO (SARS-CoV-2) or Type I/II IFN-R KO (SARS-CoV) for these studies, knock out of either or both signaling pathways may be contributing to the observed restoration of replication. Future studies are aimed at deconvoluting this issue. Nevertheless, these data suggest that SARS-CoV-2 ExoN- attenuation is not simply due to gross defects in replication but rather signals the potential for conserved CoV ExoN functions in direct and/or indirect antagonism of the innate immune response. Given the existing data in the literature on both CoV and Lassa exonuclease noted above, it is possible that SARS-CoV-2 ExoN performs one or many of the functions noted above as well as novel functions not yet reported. For example, CoV ExoN may reduce the proportion of defective viral particles or double stranded viral RNA thereby reducing agonists of the innate immune system while also potentially directly interfering with ISG protein production. Given our *in vivo* data in IFN-R KO mice, it is curious that we still find significant attenuation of SARS-CoV-2 ExoN- viral replication in our engineered Vero E6 cells (i.e. VTA), a cell type with a known genetic defect in type I IFN production (39). In the absence of a type I IFN response, Hantavirus infection of Vero E6 cells has been shown to induce type III IFN (57), which may be partially limiting SARS-CoV-2 ExoN- viruses in VTA cells. Therefore, future studies are focused on uncoupling the innate antagonist and replication fidelity phenotypes of SARS-CoV-2 ExoN- virus *in vitro and in vivo*.

Collectively, these data argue for an expansion of CoV nsp14-ExoN function. In summary, we demonstrate that CoV ExoN is not only a key mediator of replicative fitness and fidelity but is also important for subverting the innate immune response. Like our prior work with MHV and SARS-CoV, we show that SARS-CoV-2 ExoN- LOF viruses have diminished replication fidelity and competitive fitness, have increased sensitivity to nucleoside analog antivirals, have altered recombination patterns, are attenuated in primary human airway epithelial cells and are attenuated for replication and pathogenesis in typical laboratory mouse models. However, replicative fitness *in vivo* is partially restored the absence or type I/III interferon signaling indicating that *in vivo* attenuation in WT mice is not driven by gross defects in replication. These data suggest that the CoV replicase is not just a governor of viral RNA synthesis, but that nsp14-ExoN mediates virus and host interactions that are essential to subvert the cellular antiviral response. Together, these data reveal new insights into the complexities of CoV replication and virus and host interactions which could be leveraged for the development of novel multifaceted therapeutics that attack the ever expanding functions of the CoV RTC in replication and pathogenesis.

## Materials and Methods

### Cells and Viruses

#### Cells

Vero E6 cells were obtained from the United States Army Medical Research Institute of Infectious Diseases (USAMRIID) and cultured in Dulbecco’s modified Eagle medium (DMEM) (Gibco) supplemented with 10% fetal bovine serum (FBS) (Gibco), and 100 U/ml penicillin (Gibco) (Complete DMEM). Vero E6 cells overexpressing the human transmembrane protease, serine 2 (TMPRSS2) and human ACE2 receptor (VTA cells) were gifted from A. Creanga and B. Graham, National Institutes of Health (NIH) and grown in DMEM supplemented with 10% FBS and passaged in the presence of 10µg/ml puromycin dihydrochloride (Corning). Cells were routinely washed in Dulbecco’s phosphate-buffered saline without calcium chloride or magnesium chloride (PBS -/-). Cells were maintained at 37°C, 5%CO_2_, and detached during passage and expansion with 0.05% trypsin-EDTA (Gibco). BHK-21 (ATCC CCL-10) cells were propagated in DMEM (Gibco) supplemented with 10% FBS (Gibco), penicillin-streptomycin (Gibco), and L-glutamine (Gibco). MA-104 cells (ATCC CRL-2378.1) were grown in EMEM (Gibco) supplemented with 10% FBS (Gibco), penicillin-streptomycin (Gibco). Cell growth conditions during experimentation are listed in context.

#### Viruses

All infectious clone severe acute respiratory syndrome coronavirus 2 (SARS-CoV-2) were based on the WA1 isolate (WT: GenBank MT461669.1; nLuc: GenBank MT844089; MA10: GenBank MT952602) (26, 27). Wild type (WT) infectious clone SARS-CoV-2 used for this study was recovered using reverse genetics system described previously(26, 27), in Vero E6 cells at 37°C, P0 stocks. P1 stocks were generated in VTA cells at either 33°C or 37°C and used in the experiments described. SARS-CoV-2 WA1 and MA10 ExoN mutants (i.e. YQAV, RAYF, AVFS, VHVV, AAGH) were recovered through electroporation of BHK cells and coculture with VTA cells at 33°C. SARS-CoV-2 ExoN viruses were passaged twice to obtain working stocks for the studies described herein. SARS-CoV MA15 and SARS-CoV MA15 AAGH were generated as previously described (23). All working stocks were deep sequenced to identify mutations acquired during amplification. See Supplementary Table 2 and 3 for stock sequencing information.

### Saturation mutagenesis and recovery of SARS-CoV-2 nsp14-ExoN motif-I viruses

SARS-CoV-2 nsp14-ExoN motif-I mutant libraries were engineered through saturation mutagenesis on amino acid residues 90, 92, 93, and 95 of SARS-CoV-2 nsp14-ExoN motif-I based on previously published protocols (58, 59). The nsp14 gene is located in the E fragment of the SARS-CoV-2 infectious clone (27). First, a “killed” E fragment was generated to eliminate the possibility of WT contamination in the cloning process. The SARS-2 E plasmid was digested with Bsu36I and Bpu10I and replaced with a gBlock (IDT) deleting 10nt (amino acids 90-92) and inserting a BtgZI restriction site.

Degenerate NNK oligonucleotides (Integrated DNA Technologies) were used to amplify the region of interest to generate a library with mutated nsp14-ExoN motif-I DNA fragments. To limit bias and ensure accuracy, Q5 high-fidelity polymerase (NEB) was used and limited to <18 cycles of amplification. The DNA library was cloned into the SARS-CoV-2 reverse-genetics system plasmid E to create a plasmid library by Gibson cloning. The resultant E fragment library was then digested and then utilized in a ligation reaction with the remaining viral cDNA fragments to generate full-length viral cDNA. Ligation reactions were then concentrated and purified by ethanol precipitation. Purified ligation products were electroporated into DH10B ElectroMax cells (Invitrogen) and directly plated on multiple 5,245-mm2 bioassay dishes (Corning) to avoid bias from bacterial suspension cultures. Colonies were pooled and purified using a Maxiprep kit (Qiagen). The plasmid library was used for SARS-CoV-2 reverse genetics as described above. The in vitro-transcribed SARS-CoV-2 RNA library was electroporated in BHK cells and cocultured with VTA cells, and the viral supernatants were passaged two times every 4 to 5 days in VTA cells for enrichment. RNA was isolate from supernatants with TRIzol LS. Purified RNA was prepared for sequencing using the Illumina RNA Prep with Enrichment (Illumina, 20040536) workflow, following the instructions for the Respiratory Virus Oligo Panel (Illumina, 20044311). The resulting libraries were run on a MiSeq instrument, with at least 1.5 million reads per sample. Sequences were analyzed and displayed using custom Perl, Python, and R scripts as described previously(59). These scripts and usage information are available under the Saturation Mutagenesis Pipeline on the Tse lab GitHub site (https://github.com/TseLabVirology/Saturation-Mutagenesis-Pipeline/tree/main). In brief, the CAMseqv4.pl Perl script was used to extract library sequences from sequencing files. Then, these were prepared for plotting using the merge_AA.py python script to calculate amino acid distance to the DVEGCH wildtype sequence. The resulting data was input into R, enrichment scores were calculated, and plotted using bubbpleplot_enriched_AA.r.

### Mutagenesis and recovery of D90A/E92A

SARS-CoV-2 infectious clone plasmid was used as template for mutagenesis (27). Site-directed mutagenesis by “round-the-horn” PCR was used to generate substitutions at the indicated sites. SARS-CoV-2 E fragment was used as a template to mutate nucleotides A18,308C and T18,309A using the following primers: M1N-V4F (5’-CAGTCGAGGGGTGTCATGCTACTAGAGAAGCT-3’) and M1N-V2-5R (5’-CGAAGCCAATCCATGCACGTACATGTCTTATAGCTTC-3’), resulting in nsp14 amino acid substitution D90A. This plasmid was then used as a template to mutate nucleotides A18,314C and G18,315T using the following primers: M1N-V6F (5’-CTGGGTGTCATGCTACTAGAGAAGCTGTTGGTAC-3’) and M1N-V6R (5’-CGACTGCGAAGCCAATCCATGCAC-3’), resulting in the nsp14 double amino acid substitution D90A/E92A. All primers were 5’-phosphorylated with T4 polynucleotide kinase using an ATP-containing reaction buffer (NEB). Template backbone DNA was digested with DpnI (NEB), and amplified DNA was separated by electrophoresis and extracted from agarose (Promega). Ligated DNA was transformed into Top 10 competent *Eschericia coli* cells (Thermo) and amplified in liquid culture, and sequences were confirmed by Sanger sequencing. Assembly and recovery of recombinant SARS-CoV-2 have been described previously (27), with the following modifications. VTA cells were electroporated with *in vitro* transcribed, full-length genome assemblies and grown at 33°C 5%CO_2_. When the cytopathic effect consumed approximately 80% of the monolayer, clarified supernatants were aliquoted and stored at -80°C (passage 0, P0). The P0 stocks were then passaged in VTA cells at either 33°C or 37°C, P1. P1 stocks of 33°C or 37° grown virus cultures were used for most in vitro experiments presented. Engineered mutations of the P0 or P1 stocks were confirmed by Sanger Sequencing of PCR amplicons harboring the sequence of interest (each 3-4 kb in length).

### Replication kinetics of SARS-CoV-2 ExoN- viruses in VTA cells

4E+05 VTA cells were seeded per well per six well plate the previous day to generate sub-confluent monolayers. The VTA cells were infected in triplicate with the SARS-CoV-2 MA10 WT or related ExoN mutant (i.e. AAGH, RAYF, AVFS, VHVV, or YQAV) viruses at an MOI of 0.01 in Infection Media (DMEM, 5% FBS, 1X Pen/Strep) for 1hr at 37°C with rocking. After infection, the virus was removed, the monolayers were washed with 2mL PBS without Ca^2+^/Mg^2+^, and 2mL of Infection Media was added per well. The plates were incubated at 37°C, and virus supernatants were harvested at the 0 (post wash), 8, 24, 36 and 48hr post infection. At each timepoint, a 700µL sample of the infection supernatant was collected, clarified (13,000rpm, 5min), and stored at -80°C. 700µL of fresh, warmed Infection Media was added back to each well at each collection timepoint. The virus supernatants were titered by plaque assay in VTA cells as described previously (27).

### Virus replication assays to determine specific infectivity

VTA cells were plated at a density of 4E5 cells per well in 6-well plates 24h before infection. Cells were then infected at an MOI of 0.01 PFU/cell and incubated at either 33°C or 37°C for 30 minutes. Inocula were removed, and the cells were washed twice with PBS containing calcium chloride and magnesium chloride (PBS +/+). Complete DMEM was added to cells, and cells were incubated at 33°C or 37°C for 48h. At the indicated times, 700µl supernatant samples were collected, and 700µl of temperature-matched complete DMEM was added back. Of the 700µl samples collected, 100µl was added to TRIzol-LS reagent (Ambion), and the remaining 600µl was stored at -80°C for plaque assay.

### Plaque assays and RT-qPCR

Plaque assays were performed on subconfluent VTA cells seeded in 6-well plates. Serial dilutions of virus samples were plated in duplicate and overlaid with 0.9% agar in DMEM containing 5% FBS and 1x penicillin/streptomycin, cells were incubated at either 33°C or 37°, and titers were scored 72hpi. Genome quantification was determined by one-step RT-qPCR for supernatant and monolayer-derived RNAs extracted with TRIzol and purified with a KingFisher MagMAX Viral/Pathogen Nucleic Acid Isolation Kit (Thermo) according to the manufacturer’s protocol. Viral RNA was detected on a QuantStudio 3 real-time PCR system (Applied Biosystems) by TaqMan Fast Virus 1-Step Master Mix chemistry (Applied Biosystems) using a 5’ 6-carboxyfluorescein (FAM) and 3’ black hole quencher 1 (BHQ-1)-labeled probe (5’-ACCTACCTTGAAGGTTCTGTTAGAGTGGT-3’) and forward (5’-GTGCTCATGGATGGCTCTATTA-3’) and reverse (5’-TGTTGTCATCTCGCAAAGGCTCTCA-3’) primers corresponding to nsp4. RNA copy numbers were determined using an nsp4 RNA standard derived from the SARS-CoV-2 B fragment.

### Competitive fitness assay

VTA cells were plated at 4E5 cells per well in 6-well plates 24h before infection. Cells were then co-infected with three independent lineages at a total MOI of 0.01 PFU/cell at either a 1:1 ratio of WT:AAGH, MOI 0.005 PFU/cell each, or a 1:9 ratio of WT:AAGH, MOI 0.001 or 0.009 PFU/cell, respectively. Cells were incubated at either 33°C or 37° for 30 min. Inocula were removed, and the cells were washed twice with PBS +/+.

Complete DMEM, 1.5ml, was added to cells, and cells were incubated at 33°C or 37°C for 18h. The entire P1 supernatant sample was collected, and 50µl was used to blindly infect VTA cells plated at the same density 24h in advance. The entire P2 supernatant sample was collected at 18hpi. The resulting P1 and P2 cell-infected monolayers were collected in TRIzol reagent, and viral RNA was extracted by chloroform extraction and purified using the KingFisher MagMAX Viral/Pathogen Nucleic Acid Isolation Kit. Viral cDNA was generated wth SuperScript IV reverse transcriptase using random hexamers and oligo(dTs). Amplicons were generated via PCR using EasyA polymerase and Sanger sequenced. Sanger sequencing traces at genome positions 18,308; 18,309; 18,314; and 18,315 were then analyzed by area under the curve, and the pooled percentage of either wild-type or engineered mutant nucleotide were graphed.

### SARS-CoV-2 Nucleoside analog sensitivity studies

2E+04 VTA cells were seeded in 100µL Growth Media (DMEM, 10% FBS, 1X NEAA, 1X Pen/Strep, 10µg/mL Puromycin) per well in a black-bottomed 96 well plate the day prior to infection. For infection, media was aspirated and cells were infected with 100µL of SARS-CoV-2 WT or ExoN(-) AAGH nanoluciferase-expressing viruses diluted to an MOI of ∼0.1 in Infection Media (DMEM, 5% FBS, 1X Pen/Strep). The plates were incubated at 37°C for 1hr. After incubation, the virus was removed, the monolayers were washed with 100µL of Infection Media, and the cells were treated with three-fold serial dilution series of 5-fluorouracil (5-FU, Sigma), β-d-N4-Hydroxycytidine (NHC, EIDD-1931, MedChemExpress) or GS-441524 (MedChemExpress) in Infection Media, starting at concentrations of 400µM, 20µM, and 20µM respectively. The final concentration of small molecule vehicle, DMSO, was 0.2% per well. Concurrently, non-infected cells were treated with the same dose responses of the above compounds to determine cytotoxicity. The plates were incubated at 37°C for 24hr and viral replication determined through the measure of nanoluciferase activity using the NanoGlo Luciferase Assay System (Promega) and cytotoxicity was determined by CellTiterGlo Assay (Promega). Each condition was evaluated in triplicate in two independent studies. Values were normalized to the uninfected and infected vehicle DMSO controls (0 and 100% infection, respectively). Data were fit using a four-parameter nonlinear regression analysis using GraphPad Prism. EC50 and CC50 (cytotoxic concentration at which 50% of cells are viable) values were then determined as the concentration reducing the signal by 50%.

### Infections for RNA sequencing and recombination analysis

VTA cells were plated at 1E6 cells per T25 flask. Cells were infected with the indicated viruses at an MOI of 0.01and incubated at 33°C for 30 min. Inocula were removed, and the cells were washed twice with PBS +/+. Complete DMEM was added to cells, and cells were incubated at 33°C. Supernatants were removed, and the monolayers were collected in TRIzol when cells were ≈70% engaged in CPE, 20-45hpi. Viral RNA was extracted by chloroform extraction and purified using the KingFisher MagMAX Viral/Pathogen Nucleic Acid Isolation Kit. Illumina RNA sequencing of viral RNA, processing, and alignment Extracted total RNA underwent poly(A) selection followed by NovaSeq PE150 sequencing (Illumina) at 15 million reads per sample at the Vanderbilt University Medical Center core facility, Vanderbilt Technologies for Advanced Genomics (VANTAGE). VANTAGE performed base-calling and read-demultiplexing. The CoVariant pipeline was used for variant analysis (31, 35). The first module trims and aligns raw FASTQ files to the viral genome for each specified sample using a standard Bash shell script. To summarize, raw reads were processed by first removing the Illumina TruSeq adapter using Trimmomatic (60). Reads shorter than 36 bp were removed, and low-quality bases (Q score of <30) were trimmed from read ends. The raw FASTQ files were aligned to the SARS-CoV-2 genome (MT020881.1) by using the CoVariant Python 3 script and the ViReMA (Viral Recombination Mapper, version 0.21) (61) Python3 script command line parameters. For variant analysis, the sequence alignment map (SAM) file was processed using the Samtools suite (62), and alignment statistics output was generated by Samtools idxstats to an output text file. Nucleotide depth at each position was calculated from the SAM files using BBMap (Bushnell) pileup.sh.

For recombination analysis, the RecombiVIR pipeline was used (35). Following alignment, recombination junctions were filtered, quantified, and annotated by using RecombiVIR_junction_analysis.py with the following command line parameters: python RecombiVIR_junction_analysis.py samples.txt SARS2 ../directory experiment_name—version 0.21 -- Shannon Entropy ../Shannon_Entropy – Virus_Accession MT020881.1. In summary, the recombination JFreq was calculated by comparing the number of nucleotides in detected recombination junctions to the total number of mapped nucleotides in a library. JFreq was reported as the junctions per million nucleotides sequenced. Mean JFreq values are reported. Forward recombination junctions were classified as either sgmRNA junctions or DVG junctions, based on the position of their junction sites, and filtered in module 2 of RecombiVIR (RecombiVIR_junction_analysis.py). Briefly, junction start sites were filtered to those positioned within 30 nucleotides of the transcriptional regulatory sequence leader (TRS-L) for each virus. The stop sites were then filtered for those positioned within 30 nucleotides of each respective sgmRNA TRS. This window is supported by other reports defining the flexibility of the CoV transcriptome. The JFreq of each sgmRNA was calculated by dividing the number of nucleotides in a specific sgmRNA population by the total amount of viral RNA (total mapped nucleotides). This ratio was multiplied by 10^6^ to scale or the number of nucleotides sequenced. DVG JFreq was calculated by dividing the number of nucleotides in DVG junctions by the total amount of viral RNA in a sample (total mapped nucleotides). This ratio was multiplied by 10^6^ to scale or the number of nucleotides sequenced. The percentage of sgmRNA and DVG junctions was then calculated by comparing the depth of all filtered sgmRNA or DVG junctions to the sum of all detected forward junctions.

### Infections for RNA sequencing of genomic RNA from DMSO or 5’FU treated cells

T-25 flasks of VTA cells seeded the day prior with 1E+06 cells were infected with SARS-CoV-2 WT, AAGH or YQAV at an MOI of 0.01 for 1hr at 33°C after which input virus was removed, monolayers were washed with PBS and Infection Media was added including DMSO vehicle or 100µM 5’-FU. After 24hr, clarified supernatants were harvested, RNA extracted in TRIzol LS and analyzed by Illumina MiSeq. Each condition was performed in quadruplicate. The total number of variants, the number of variants > 1% and <1% normalized to the number of reads mapped to the reference sequence

### SARS-CoV-2 nsp10/nsp14 fusion cloning protein expression

SARS-CoV-2 NSP10 fused via a 2X GGS linker to NSP14 was codon optimized and cloned into vector pK27, which includes three N-terminal tags: his tag, flag tag, and a SUMO tag. C41 (DE3) pLysS competent cells (Sigma-Aldrich CMC0018) were transformed and grown in terrific broth at 37°C with 50 µg/mL kanamycin until the cultures reached a density between 0.8 and 1 OD600. Cultures were cooled down at 4°C for 1 hour, induced with 0.5 mM isopropyl β-D-1-thiogalactopyranoside (IPTG), and incubated overnight on a shaker at 200 RPM and 18°C. Next day, the cultures were centrifuged at 4,000 xg for 10 minutes, and the cell pellets were resuspended in 20 mL of lysis buffer (50 mM NaH2PO4, 300 mM NaCl, 10 mM imidazole, pH 8, complete EDTA-free protease inhibitor tablet (Sigma-Aldrich 04693132001)) per 2-5 g cell pellet weight, then lysed with a cell dismembrator. Lysed cells were then centrifuged at 18,000 xg for 30 minutes at 4°C, and the supernatant was passed through a 0.45 µm filter and resuspended in a ratio of 1:5 in lysis buffer and incubated with Talon affinity resin (Takara) at 4°C for 1 hour, with occasional shaking. Resin was captured via gravity chromatography, and the protein removed with elution buffer (50 mM NaH2PO4, 300 mN NaCl, and 500 mM imidazole pH 8), then dialyzed overnight twice against PBS to remove imidazole. Protein was then concentrated with a PES concentrator (Thermo Sci., 88541), quantitated via protein assay kit (Thermo Sci., A53225), and visualized with a 4-20% mini-Protean TGX Stain-free gel (BioRad) imaged with a BioRad Chemidoc MP imaging system.

### FRET-based *in vitro* exonuclease assay

Exonuclease activity was evaluated using purified WT and mutant NSP10/NSP14 proteins diluted in NSP buffer (50 mM Tris-HCl, 0.0001% [vol/vol] Tween 20, 5% [vol/vol] glycerol, 1.5 mM MgCl2, 20 mM NaCl, 0.5 mM tris(2-carboxyethyl)phosphine [TCEP], 0.1 μg/mL bovine serum albumin [BSA]). Proteins were left with the affinity tags intact. 500 nM WT protein was mixed with 250 nM annealed RNA oligos containing fluorophore and quencher 5′TexRd-XN/rArCrArArArArCrGrGrCrCrCrA and rA*rA*rA*rU*rA*rG*rG*rG*rC*rC*rG*rU*rU*rU*rU*rG*rU*/3′IAbRQSp/ (* designates phosphothioate bond). Oligos were combined 1:1 and annealed starting at 95°C for 10 minutes, with a 5°C stepdown every minute thereafter until a final 5-minute step at 25°C. Fluorescence was measured using a plate reader (Thermo Sci., Varioskan Lux 3020–81011), at an excitation of 590 nm, and emission 615 nm for 1hr at room temperature. Interface and adaptive mutant proteins were normalized to WT concentrations using a combination of protein quantitation assay results and protein band intensity imaged in a mini-Protean TGX Stain-Free gel.

### Evaluating the susceptibility of SARS-CoV and SARS-CoV-2 ExoN- viruses to IFNβ pretreatment in vitro

For <u>SARS-CoV-2</u>, 2E+04 VTA cells were seeded in 100µL of Growth Media (DMEM, 10% FBS, 1X NEAA, 1X Pen/Strep, 10µg/mL Puromycin) per well in a black-bottomed 96 well plate. The following day, the VTA cells were pre-treated with a three-fold serial dilution series of IFNβ, starting with a top concentration of 10ng/mL, prepared in Growth Media (DMEM, 10% FBS, 1X NEAA, 1X Pen/Strep, 10µg/mL Puromycin) for 24hrs. The IFNβ-containing growth media was removed and the VTA cells were infected with 100µL of the SARS-CoV-2 WT or ExoN(-) AAGH nanoluciferase-expressing viruses diluted to an MOI of ∼0.1 in Infection Media (DMEM, 5% FBS, 1X Pen/Strep). The plates were incubated at 37°C for 1hr. After incubation, the virus was removed, the monolayers were washed with 100µL of Infection Media, and 100µL of Infection Media was added to each well. The plates were incubated at 37°C and nanoluciferase activity (used as a proxy for virus replication) was measured at 72hpi using the NanoGlo Luciferase Assay System (Promega). For <u>SARS-CoV</u>, 1E+04 MA-104 cells/well in a 48 well plate were seeded 48hr prior to infection. 6hr prior to infection, cells were exposed to a dose response of IFN-ß (InvivoGen): 0, 10, 100, or 500 units/mL. Cells were then infected with SARS-CoV MA15 or SARS-CoV MA15 AAGH at an MOI of 1 assuming cells have doubled. After 48hr, the levels of infectious virus in the supernatant were measured via plaque assay.

### Replication kinetics of SARS-CoV-2 ExoN- viruses in human airway epithelial cell cultures (HAEs)

HAE cell cultures from two independent human donors were obtained from the Tissue Procurement and Cell Culture Core Laboratory in the Marsico Lung Institute/Cystic Fibrosis Research Center at UNC. Cells were maintained in “air liquid interface” (ALI, Tissue Procurement and Cell Culture Core Laboratory) medium. Prior to infection, the apical side of each culture was washed with 0.5mL of PBS without Ca^2+^/Mg^2+^ for 30min at 37°C to remove excess mucus. For infection, 200µL of SARS-CoV-2 MA10 WT or related ExoN mutant AAGH, or YQAV viruses (MOI 0.1) was added to the apical surface of HAE in triplicate and incubated 1.5hrs at 37°C after which input virus was removed and the apical surfaces were washed with 0.5mL Ca^2+^/Mg^2+^ for 10min at 37°C to wash away input virus. To determine infectious virus production, the apical surfaces of cultures were washed with 200µL PBS after 24, 48, and 72hr and titered by plaque assay in VTA cells as described above.

### Assessing replication and pathogenesis of SARS-CoV-2 ExoN- viruses in Balb/c mice

10-week-old, female Balb/c mice were anesthetized with ketamine-xylazine and infected intranasally with 6.8E+05 PFU of WT SARS-CoV-2 MA10 (N = 15) or related YQAV virus (N = 15) diluted in PBS without Ca^2+^/Mg^2+^. Body weights were measured daily. On 1, 2 and 4dpi, a subset of animals (N = 5/group) were humanely euthanized, gross lung pathology score was recorded and the inferior right lung lobe was harvested and stored at -80°C until infectious virus titration by plaque assay in VTA cells as described previously(26). Gross lung pathology is observed with emerging CoV infection of mice where lungs can appear “hemorrhaged”. This gross lung pathology phenotype is scored on a scale of 0–4, where 0 is a normal pink healthy lung and 4 is a completely dark red lung.

### Assessing replication and pathogenesis of SARS-CoV-2 ExoN- viruses in in K18-hACE2 mice

8-9 month old male and female K18-hACE2 mice were anesthetized with ketamine-xylazine and infected intranasally with 3E+04 PFU WT SARS-CoV-2 MA10 (N = 12) or related ExoN- viruses AAGH (N = 12), RAYF (N = 10), AVFS (N = 10), VHVV (N = 10), or YQAV (N = 12) diluted in PBS without Ca^2+^/Mg^2+^. Body weights were measured daily. On 2dpi, 4 animals per group were humanely euthanized, gross lung pathology score was recorded and the inferior right lung lobe was harvested and stored at -80°C until infectious virus titration by plaque assay in VTA cells as described previously (26). On 6dpi, the remaining animals were humanely euthanized, gross lung pathology score was recorded, the inferior right lung lobe and the left brain hemisphere was harvested and stored at -80°C until infectious virus titration by plaque assay in VTA cells.

### Assessing replication and pathogenesis of SARS-CoV-2 ExoN- viruses in type I (αβ) and type III (λ) IFN receptor double knockout (DKO) B6 mice

8-15 week old male and female WT C57BL/6J (Jackson Labs) and congenic C57BL/6J IFN αβλ-R KO mice were anesthetized with ketamine-xylazine and infected intranasally with 6.8E+05 PFU of WT SARS-CoV-2 MA10 (WT N = 10, KO N = 11) or related YQAV virus (WT N = 10, KO = 15) diluted in PBS without Ca^2+^/Mg^2+^. Weights were measured daily, and on 4dpi, animals were humanely euthanized, gross lung pathology score was recorded and the inferior right lung lobe was harvested and stored at -80°C until infectious virus titration by plaque assay in VTA cells as described previously (26).

### Assessing replication and pathogenesis of SARS-CoV MA15 ExoN- viruses in type I (αβ) and type II (γ) IFN receptor double knockout (DKO) B6 mice

20 week old female WT C57BL/6J (Jackson Labs) and congenic C57BL/6J IFN αβγ-R KO mice were anesthetized with ketamine-xylazine and infected intranasally with 6.8E+05 PFU of WT SARS-CoV MA15 (WT N = 27, KO N = 25) or related SARS-CoV MA15 AAGH virus (WT N = 42, KO = 41) diluted in PBS without Ca^2+^/Mg^2+^. Weights were measured daily. On 1, 2 and 4dpi, a subset of animals from each group were humanely euthanized and the inferior right lung lobe was harvested and stored at -80°C until infectious virus titration by plaque assay in Vero-E6 cells as described previously (63).

### Laboratory Biosafety

All in vitro and in vivo work presented herein was approved by either the Vanderbilt University, the UNC Institutional Biosafety Committee and or the Institutional Animal Care and Use Committee at UNC Chapel Hill. All virology was performed with approved standard operating procedures for SARS-CoV or SARS-CoV-2 in BSL3 facilities which met requirements recommended in “Biosafety in microbiological and biomedical laboratories” by the US Department of Health and Human Service, the US Public Health Service, the US Center for Disease Control and Prevention, and the NIH.

### Statistical analysis

GraphPad Prism, version 9 (La Jolla, CA) was used for all statistical analyses. All tests and sample sizes are listed in the figure legends.

## Data availability

FASTQ files for the RNA sequencing variant and recombination analyses have been deposited in the National Center for Biotechnology Information Sequence Read Archive under the accession numbers: PRJNA1244841, PRJNA1245334. Growth of virus in the presence of small molecule treatment FASTQ files can be found in Sequence Read Archive under the accession number PRJNA1374231. FASTQ files for the sequencing of saturation mutagenesis viral supernatants have been deposited in the Sequence Read Archive under the accession number PRJNA1391540.

---

**Figure S1: SARS-CoV-2 ExoN inactivating mutations abolish enzymatic activity.** The *in vitro* exonuclease activity for select ExoN mutants was determined by a fluorescence RNA cleavage assay using nsp10/nsp14 fusion proteins. The RNA cleavage activity of WT and mutant nsp14 fusion proteins was measured over time using normalized amounts of protein and double stranded RNA FRET robe. Symbols represent mean values and ± SEM error bars from four independent experiments.

**Figure S2: SARS-CoV-2 AA has altered recombination.** Associated with main Figure 5. VTA-infected monolayer RNAs from experiments shown in Figure 5 were analyzed by RNAseq and ViReMA. **(A)** The junction frequency (JFreq) was calculated as the ratio of detected junctions per 1 million mapped nucleotides. **(B)** Junction frequencies were calculated for defective viral genomes (DVGs) and total subgenomic RNAs (sgmRNA) and plotted as the percentage of total mapped junctions. **(C)** Individual sgmRNA junction frequencies from panel B are shown as the percentage of total mapped junctions. Graphed are individual values, mean, and ± SEM error bars from three independent experiments, n=3. **p<0.01, ***p<0.001, ****p<0.0001, ns not significant as determined by unpaired t-test. WT **(D)** and AA (E) recombination junctions are mapped according to their genomic position (5’ junction site starting position, 3’ junction site sop position) and colored according to their frequency in the population of all mapped junctions. The highest frequencies are magenta, and the lowest frequencies are red. Dashed boxes represent clusters of junctions (i) 5’→ 3’, (ii) mid genome → 3’, (iii) 3’ → 3’, (iv) local deletions, (v) 5’ UTR → rest of genome. Shown are representative results from one of three independent experiments with similar outcomes.

**Figure S3: Interferon deficiency partially restores SARS-CoV ExoN- replicative fitness in vivo.** Associated with Main Figure **(A)** SARS-CoV MA15 and AAGH virus interferon beta sensitivity. MA-104 cells were pretreated with a dose response of interferon beta 6hr prior to infection with an MOI of 1. Infectious virus production was measured at 48hpi by plaque assay. **(B)** Schematic of in vivo pathogenesis study in WT C57BL/6 or interferon alpha, beta, gamma receptor knock out mice infected with WT SARS-CoV MA15 (WT N = 27, KO N = 25) or SARS-CoV MA15 AAGH (WT N = 42, KO = 41). **(C)** Body weight loss over time for mice described in “B”. The symbol represents the mean and the error bars represent the standard deviation. **(D)** Viral lung titers for mice described in “B”. The line is at the mean and the error bars represent the standard error of the mean. Asterisks in C and D indicate statistical significance by Two-way ANOVA with Sidak’s multiple comparison test.

**Table S1: Sequences of viable saturation mutagenesis variants enriched in passage.** The amino sequence from residue 90-95, the amino acid differences from WT DVEGCH sequence, the percentage in the population and the enrichment score is shown. The enrichment score was generated by comparing the percentage of a given variant at P0 to the percentage of that same variant after 2 passages.

**Table S2: Coding mutations identified in SARS-CoV-2 AAGH virus stocks.** RNA was extracted from cell monolayers that produced the indicated viral stocks and analyzed by RNAseq. Non-engineered variants are listed that met the following conditions: i) at least 10% frequency, ii) coded for amino acid substitution, iii) not identified in WT stock. Variants that did not meet the 10% frequency cut off were included if they met the conditions listed above for at least one virus stock.

**Table S3: Coding mutations identified in SARS-CoV-2 MA10 nsp14 active site virus stocks.** RNA was extracted from cell monolayers that produced the indicated viral stocks and analyzed by RNAseq. Non-engineered variants are listed that met the following conditions: i) at least 10% frequency, ii) coded for amino acid substitution, iii) not identified in WT stock. Variants that did not meet the 10% frequency cut off were included if they met the conditions listed above for at least one virus stock.

